# Preservation of Mitochondrial Membrane Potential is Necessary for Lifespan Extension from Dietary Restriction

**DOI:** 10.1101/2022.12.27.522028

**Authors:** Brandon J. Berry, Evan Mjelde, Fatima Carreno, Kathryn Gilham, Emily J. Hanson, Emily Na, Matt Kaeberlein

## Abstract

Dietary restriction (DR) increases lifespan in many organisms, but its underlying mechanisms are not fully understood. Mitochondria play a central role in metabolic regulation and are known to undergo changes in structure and function in response to DR. Mitochondrial membrane potential (Δψ_m_) is the driving force for ATP production and mitochondrial outputs that integrate many cellular signals. One such signal regulated by Δψ_m_ is nutrient-status sensing. Here, we tested the hypothesis that DR promotes longevity through preserved Δψ_m_ during adulthood. Using the nematode *Caenorhabditis elegans*, we find that Δψ_m_ declines with age relatively early in the lifespan, and this decline is attenuated by DR. Pharmacologic depletion of Δψ_m_ blocked the longevity and health benefits of DR. Genetic perturbation of Δψ_m_ and mitochondrial ATP availability similarly prevented lifespan extension from DR. Taken together, this study provides further evidence that appropriate regulation of Δψ_m_ is a critical factor for health and longevity in response to DR.

## INTRODUCTION

Dietary restriction (DR) can be defined as decreased nutrient availability in the absence of malnutrition. A variety of different DR regimens have been shown to extend lifespan across taxa, from single-celled budding yeast, to invertebrate models such as nematode worms and fruit flies, to rodents, to primates [1-4]. In the nematode worm, *Caenorhabditis elegans*, DR can be implemented by complete removal from the bacterial food source during adulthood, referred to as bacterial deprivation [5, 6], by dilution of the bacterial food [7], or by mutations that reduce food consumption, collectively referred to as *eat* mutants [8]. DR in the worm is known to engage a complex nutrient-sensing network that regulates metabolic function, growth and reproduction, stress resistance, and longevity. Key components of this network include the insulin/IGF-1-like signaling pathway, xenobiotic resistance factors, and energy sensors such as AMP-activated protein kinase (AMPK) and the mechanistic target of rapamycin (mTOR) [2].

Numerous studies have suggested that mitochondrial function plays a central role in longevity determination and the response to DR [9-11], yet the mechanisms underlying this relationship remain unclear [12]. Mitochondrial membrane potential (Δψ_m_) is the voltage potential of the electrochemical proton gradient across the mitochondrial inner membrane that powers ATP production. In yeast, an early indicator of cellular aging is loss of Δψ_m_ [13]. Δψ_m_ declines with early age similarly in worms [14] and other animals [15-18]. We recently reported that increasing Δψ_m_ is sufficient to extend healthy lifespan in *C. elegans* [14]. This led us to consider how DR may affect Δψ_m_ specifically, and if changes in Δψ_m_ causally affect longevity from DR in worms.

To test the hypothesis that DR functions through preservation of Δψ_m_ with age, we performed DR on animals while perturbing Δψ_m_. Using both genetic and pharmacologic approaches, we observed that DR is sufficient to attenuate the age-associated loss of Δψ_m_ and that this is necessary for lifespan extension. These findings support a central role for mitochondrial function in the longevity-promoting effects of DR and emphasize the need for a deeper understanding of *in vivo* bioenergetics in the context of healthy longevity.

## METHODS

### C. elegans *maintenance*

Strains used in this study were the N2-Bristol wildtype strain, the DA465 strain harboring an *eat-2* mutation, the CY121 strain harboring the *upc-4* mutation, the VC620 strain harboring the *ant-1*.*2* mutation, each obtained through the Caenorhabditis Genetics Center (CGC, NIH P40 OD010440). Strain RN70 harboring the *mai-2* mutation was provided by the laboratory of Dr. Rosa E. Navarro González. We used standard nematode culture methods [19] including nematode growth media (NGM) plates and OP50 culture bacteria as a food source. Liquid media was M9 buffer (22 mM KH2PO4, 42 mM Na2HPO4, 86 mM NaCl, 1 mM MgSO4, pH 7).

### Fluorescence analysis

TMRE was dissolved in DMSO and worms were exposed at a final concentration of 1 μM in M9 buffer (0.0001% DMSO final concentration) for 24 hours while rotating. Animals stained with TMRE were exposed to FCCP in 500 μL M9 at a final concentration of 10 μM for four hours. Perhexiline was used at 30 μM final concentration for 24 hours according to previous protocols [20, 21]. MitoTracker Green FM was dissolved in DMSO and worms were exposed at a final concentration of 10 μM in 500 μL M9 buffer for 24 hours. For imaging, worms were anesthetized in 4 mM tetramisole in M9 mounted on 2% agarose pads. Fluorescence was captured on a Zeiss SteREO Lumar.V12 stereoscope (Thornwood, NY, USA). Mean and maximum fluorescence of the pharyngeal tissue was quantified using ROIs in ImageJ.

### Lifespan measurement

Age-synchronized populations were obtained by egg lay. At late L4 stage, all animals were transferred to culture plates seeded with OP50 bacterial food and 50 μM 5-fluorodeoxyuridine (FUDR) to prevent progeny hatching and development. Animals were kept at 20° C and were scored at least every other day by gently touching the head. Animals that did not move in response to touch were scored dead and removed from the plate. At day 2 of adulthood, dietary restriction (DR) populations were transferred to culture plates containing 50 μM FUDR without food as previously described [5, 6, 22] according to bacterial deprivation protocol. Also at day 2 of adulthood, populations were transferred to NGM plates containing 10 μM FCCP. Control populations were simultaneously exposed to vehicle (0.001% ethanol). Fed animals were transferred to new plates as needed to replenish food. Some of the DR animals would display foraging behavior and leave the plate, however, foraging loses living animals, therefore under-representing increase in lifespan in response to DR [5]. All lifespans were from at least 3 biological replicates (separate broods) across at least three technical replicates.

### Mobility measurement

Body bends were counted for 30 seconds while worms thrashed in M9 as previously described [23-25]. Resultant body bends per 30 seconds was multiplied by 2 to present body bends per minute in liquid for all conditions.

### Statistics

Survival curves were analyzed by Log Rank (Mantel-Cox) test. Two groups were compared using two sample two-tailed unpaired t-tests. More than two groups were compared by ANOVA with appropriate post-hoc tests. Statistical details are outlined in the figure legends for each experiment and in Supplementary table 1 for lifespans. All tests were carried out in GraphPad Prism 9.3.0.

## RESULTS

### Dietary restriction (DR) increases mitochondrial membrane potential (Δψ_m_) *in vivo*

We first confirmed that *in vivo* Δψ_m_ was decreased by day 4 of adulthood [14] by measuring TMRE fluorescence intensity in the mitochondria-rich pharynx (Figure 1A) [26, 27]. DR by bacterial deprivation (BD) significantly increased Δψ_m_ compared to the fed control (Figure 1B). Mitochondrial mass was unchanged by this DR paradigm (Supplementary Figure 1A). Similar results were obtained by measuring maximum TMRE fluorescence to avoid potential artifacts from differences in mitochondrial content (Supplementary Figure 1B) [26-30]. DR is reported to increase mitochondrial mass in some models under some conditions [31, 32], however, in other cases mitochondrial mass remains unchanged by DR [33-35], in line with our results.

**Figure 1.**
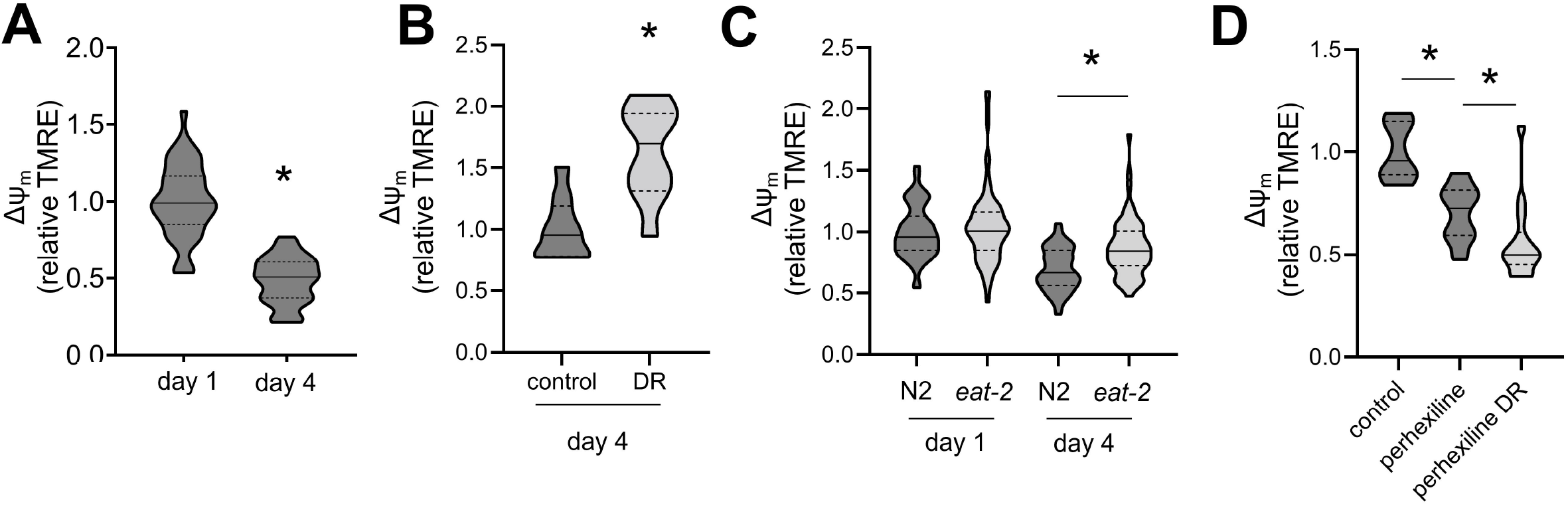
Dietary restriction (DR) causes increased mitochondrial membrane potential (Δψ_m_) A) Relative TMRE fluorescence in day 1 adult animals versus day 4 adults. Unpaired two-tailed t test, *p < 0.0001. N = 45 animals for day 1, 41 animals for day 2. Violin plots are medians ± quartiles (dotted lines). B) Relative TMRE fluorescence in day 4 adult animals either fully fed or subjected to dietary restriction (DR) by bacterial deprivation. Unpaired two-tailed t test, *p = 0.0003. N = 10 animals each condition. Violin plots are medians ± quartiles (dotted lines). Throughout, dark gray represents fed animals, and light gray represents DR. C) Relative TMRE fluorescence in day 1 adult animals versus day 4 adults in both wildtype (N2) and genetic DR (eat-2) animals. One-way ANOVA with Tukey’s multiple comparisons test, *p < 0.0001. N2 day 1 vs eat-2 day 1 p = 0.18. All other comparisons p < 0.0001. N2 day 1 N = 47, N2 day 4 N = 30, eat-2 day 1 N = 85, eat-2 day 4 N = 43. Violin plots are medians ± quartiles (dotted lines). D) Relative TMRE fluorescence in day 4 adult animals that were fully fed, subjected to DR, and treated with 30 μM perhexiline as indicated. One-way ANOVA with Tukey’s multiple comparisons test, control vs perhexiline *p < 0.0001, perhexiline vs perhexiline DR * p = 0.0409, control vs perhexiline DR p < 0.0001. Control N = 10, perhexiline N = 17, perhexiline DR N = 14. Violin plots are medians ± quartiles (dotted lines).

In order to determine whether preservation of Δψ_m_ during aging is unique to the BD method of DR, we repeated these experiments using the *eat-2* genetic model of DR. Animals with loss-of-function in the *eat-2* gene consume less food than wildtype due to a defect in pharyngeal pumping and have extended lifespan [8]. We again observed increased Δψ_m_ in the *eat-2* animals compared to wildtype controls at day 4 of adulthood (Figure 1C). Together these results suggest that DR results in preserved Δψ_m_ during aging.

There is emerging evidence that fat metabolism and mitochondrial function play a fundamental role in DR longevity in both *C. elegans* [21] and mammals [36]. BD upregulates fatty acid oxidation by mitochondria, a process that is required for DR-mediated longevity [21]. Therefore we tested the hypothesis that fatty acid oxidation is what fuels preservation of Δψ_m_ by DR by directly inhibiting fat oxidation with the drug perhexiline [20]. Prior work has shown that perhexiline prevents lifespan extension from DR [21], and we found that perhexiline prevented preservation of Δψ_m_ in response to DR and, instead, significantly reduced it (Figure 1D). Perhexiline also significantly reduced Δψ_m_ compared to fed control animals, suggesting fatty acid oxidation supports baseline *in vivo* Δψ_m_.

### Preservation of Δψ_m_ is required for DR-mediated longevity

To test the model that preserved Δψ_m_ by DR is required for longevity, we exposed animals to the mitochondrial uncoupler, FCCP, and measured the effect on lifespan under control and BD conditions. FCCP exposure was initiated at the same time as BD, day 2 of adulthood. We confirmed that FCCP decreased Δψ_m_ *in vivo*, as expected (Figure 2A), did not affect mitochondrial mass (Supplementary Figure 1C), and that FCCP exposure entirely prevented lifespan extension from DR (Figure 2B). Importantly, we chose a dose of FCCP that did not affect wildtype lifespan (Figure 2B&C). FCCP treatment similarly prevented lifespan extension by *eat-2* mutation (Figure 2C).

**Figure 2.**
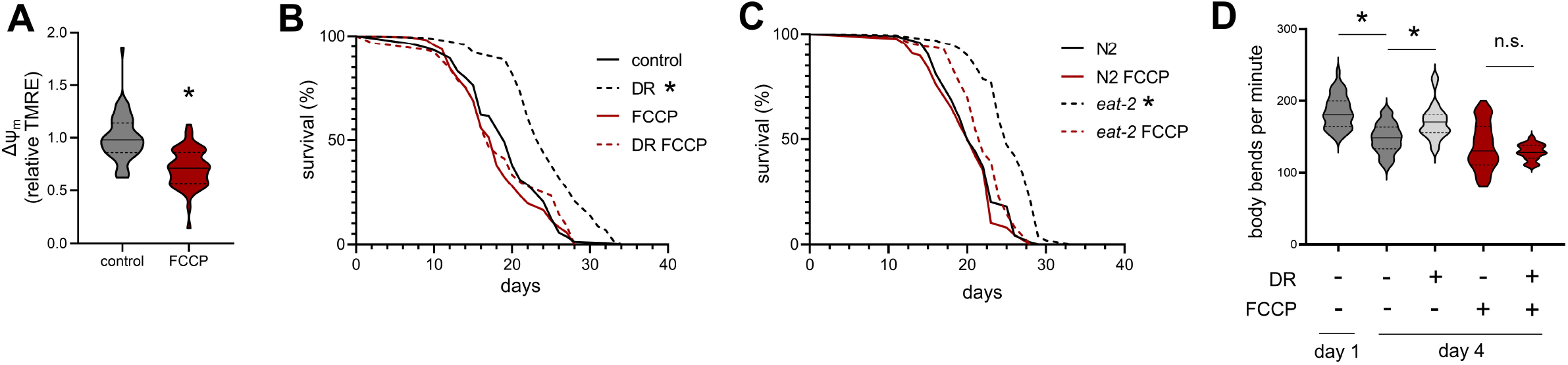
DR-mediated longevity is sensitive to decreased Δψ_m_. A) Relative TMRE fluorescence in day 4 adult animals treated with either vehicle control or 10 μM FCCP. Unpaired two-tailed t test, *p < 0.0001. Control N = 57, FCCP N = 56. Violin plots are medians ± quartiles (dotted lines). Throughout, FCCP treatment is denoted by dark red. B) Survival curves of wildtype animals (N2) subjected to DR and treated with FCCP. Log-rank (Mantel-Cox) test, *p < 0.0001. Detailed statistical information for all lifespans is presented in Supplementary Table 1. C) Survival curves of wildtype (N2) and genetic DR (eat-2) animals treated with FCCP. Log-rank (Mantel-Cox) test, *p < 0.0001. Detailed statistical information for all lifespans is presented in Supplementary Table 1. D) Motility of wildtype day 4 adult animals scored by counting body bends per minute while thrashing in liquid. Animals were subjected to DR and FCCP treatment for 2 days before measurement. One-way ANOVA with Tukey’s multiple comparisons test, N2 day 1 vs N2 DR day 4 p = 0.0393, N2 day 4 vs N2 day 4 FCCP p = 0.1270, N2 day 4 vs N2 day 4 DR FCCP p = 0.0031, N2 day 4 vs N2 day 4 DR FCCP p = 0.5545, all other comparisons p < 0.0001. N2 day 1 N = 40 animals, N2 day 4 N = 45, N2 day 4 DR N = 49, N2 day 4 FCCP N = 70, N2 day 4 DR FCCP N = 57. Violin plots are medians ± quartiles (dotted lines).

To assess whether preservation of Δψ_m_ also mediated healthspan benefits from DR, we quantified animal motility via thrashing in liquid. We found that by day 4 of adulthood control animals were significantly motility-impaired, and BD was able to reverse the impairment (Figure 2D). Upon treatment with FCCP we found that BD was no longer able to rescue age-associated loss of motility (Figure 2D).

### DR increases Δψ_m_ in *ucp-4* mutants, but not in *ant-1*.*2* or *mai-2* mutants

To begin to characterize the mechanism of increased Δψ_m_ in response to DR, we assessed Δψ_m_ of animals with loss-of-function in three genes that endogenously regulate Δψ_m_ in different ways (Figure 3A). We first tested if animals with non-functional uncoupling protein (UCP) responded to DR. UCPs result in decreased Δψ_m_ when activated [37, 38]. Therefore, we hypothesized that DR may inhibit UCP and result in increased Δψ_m_. The only known UCP in *C. elegans* is encoded by the *ucp-4* gene [39, 40]. As expected, mutation *of ucp-4* results in animals with increased Δψ_m_ at day 1 of adulthood (Figure 3B). Unexpectedly, Δψ_m_ in the *ucp-4* mutant decreased to the same level as wildtype animals by day 4 of adulthood (Figure 3B) and *ucp-4* mutants showed increased Δψ_m_ in response to DR (Figure 3C).

**Figure 3.**
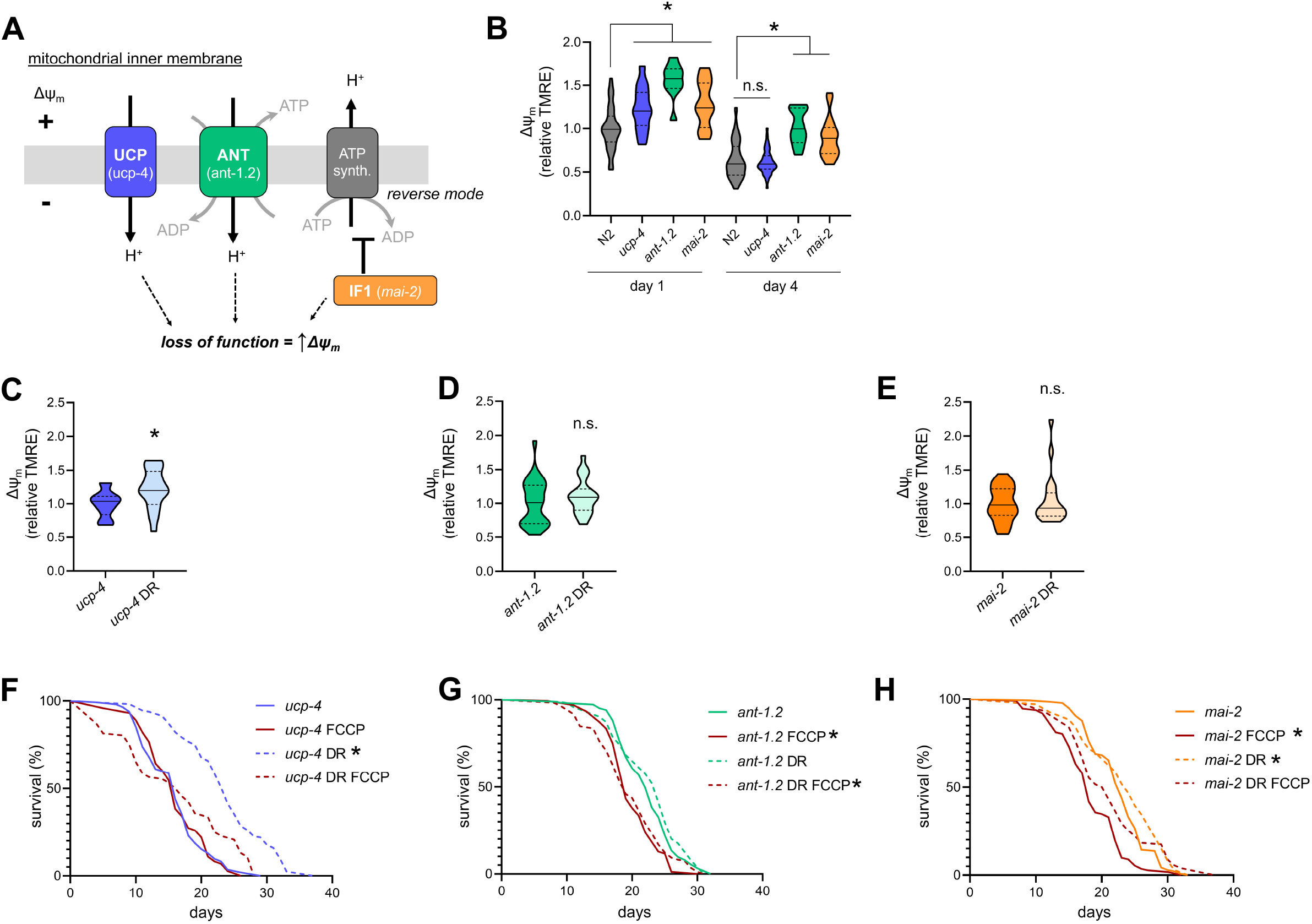
ANT and IF1, but not UCP, mediate the effects of DR in C. elegans. A) Schematic showing the mitochondrial inner membrane and proteins that regulate Δψ_m_. Uncoupling proteins (UCP) allow proton (H^+^) reentry to the mitochondrial matrix, dissipating Δψ_m_. C. elegans has a single ucp-4 gene. Adenine Nucleotide Translocase (ANT) transports ADP and ATP into and out of mitochondria, respectively, and allow proton reentry to the matrix. C. elegans ant-1.2 is highly expressed in pharynx. Inhibitory factor 1 (IF1) prevents ATP synthase from working in reverse mode, thereby preventing increased Δψ_m_. C. elegans IF1 is encoded by the mai-2 gene. Loss of function in each of these genes should result in increased Δψ_m_. B) Relative TMRE fluorescence in both day 1 adult and day 4 adult animals for wildtype (N2), UCP loss of function mutants (ucp-4), ANT loss of function mutants (ant-1.2) and IF1 loss of function mutants (mai-2). Within day 1, One-way ANOVA with Dunnett’s multiple comparisons test versus N2, ucp-4 p = 0.0007, ant-1.2 p < 0.0001, mai-2 p = 0.0006. N2 N = 40, ucp-4 N = 28, ant-1.2 N = 10, mai-2 N = 16. Within day 4, One-way ANOVA with Dunnett’s multiple comparisons test versus N2, ucp-4 p = 0.8209, ant-1.2 p < 0.0001, mai-2 p < 0.0001. N2 N = 63, ucp-4 N = 48, ant-1.2 N = 53, mai-2 N = 56. Violin plots are medians ± quartiles (dotted lines). C) Relative TMRE fluorescence in ucp-4 day 4 adult animals either fully fed or subjected to DR. Unpaired two-tailed t test, *p < 0.0218. Control N = 14, DR N = 19. Violin plots are medians ± quartiles (dotted lines). D) Relative TMRE fluorescence in ant-1.2 day 4 adult animals either fully fed or subjected to DR. Unpaired two-tailed t test, p = 0.3014. Control N = 26, DR N = 23. Violin plots are medians ± quartiles (dotted lines). E) Relative TMRE fluorescence in mai-2 day 4 adult animals either fully fed or subjected to DR. Unpaired two-tailed t test, p = 0.5686. Control N = 23, DR N = 19. Violin plots are medians ± quartiles (dotted lines). F) Survival curves of ucp-4 animals subjected to DR and treated with FCCP. Log-rank (Mantel-Cox) test, *p < 0.0001. Detailed statistical information for all lifespans is presented in Supplementary Table 1. G) Survival curves of ant-1.2 animals subjected to DR and treated with FCCP. Log-rank (Mantel-Cox) test, *p < 0.0001. Detailed statistical information for all lifespans is presented in Supplementary Table 1. H) Survival curves of mai-2 animals subjected to DR and treated with FCCP. Log-rank (Mantel-Cox) test, control vs DR *p = 0.0072, control vs FCCP *p < 0.0001. Detailed statistical information for all lifespans is presented in Supplementary Table 1.

We next tested whether the adenine nucleotide transferase (ANT) plays a role in increased Δψ_m_ in response to DR. ANT functions to transport ATP out of mitochondria and cytosolic ADP into mitochondria, and can also result in mitochondrial uncoupling similar to UCPs [41, 42] (Figure 3A). Therefore we hypothesized that DR may act through ANT activity to modulate Δψ_m_. Worms with loss of function in the *ant-1*.*2* gene showed increased Δψ_m_ at day 1 of adulthood (Figure 3B), as expected [43]. Conversely to *ucp-4*, however, *ant-1*.*2* mutants had increased Δψ_m_ compared to wildtype at day 4 of adulthood (Figure 3B), indicating that Δψ_m_ decline with age was attenuated in these mutants. When subjected to BD, Δψ_m_ was not increased in *ant-1*.*2* mutants (Figure 3D), a departure from the effect of BD in wildtype and in *ucp-4* animals.

We last tested the effects of DR on Δψ_m_ in animals defective for the mammalian inhibitory factor 1 (IF1), or *mai-2* in worms. IF1 normally inhibits ATP synthase reversal, which prevents ATP synthase from hydrolyzing mitochondrial ATP to pump protons out of mitochondria to increase Δψ_m_ (Figure 3A). ATP synthase reversal is well-described under conditions of energetic crisis [44], and IF1 is normally active to prevent this reversal [45]. Therefore, inhibiting IF1 using the *mai-2* mutation resulted in increased Δψ_m_ at day 1 of adulthood (Figure 3B), as expected [46]. Similar to animals defective for *ant-1*.*2, mai-2* animals also had increased Δψ_m_ at day 4 of adulthood (Figure 3B), suggesting that Δψ_m_ decline with age is attenuated. When subjected to BD, *mai-2* mutants did not show increased Δψ_m_ (Figure 3E) similar to the *ant-1*.*2* animals.

### DR extends lifespan of *ucp-4* mutant worms but not of *ant-1*.*2* or *mai-2* mutants

Based on their impact on Δψ_m,_ we predicted that the *ucp-4* would respond normally to DR, while the *ant-1*.*2* and *mai-2* mutant strains may experience reduced lifespan extension following DR. Consistent with this, *ucp-4* animals had increased lifespan in response to DR which was reversed by FCCP treatment, similar to wildtype animals (Figure 3F). Also similar to wildtype, FCCP had no effect on lifespan of fully fed *ucp-4* animals. In contrast, both the *ant-1*.*2* and *mai-2* strains did not experience full lifespan extension in response to DR (Figure 3G&H, Supplementary Table 1). When exposed to FCCP, *ant-1*.*2* mutants and *mai-2* mutants both experienced decreased lifespan. This sensitivity to mitochondrial uncoupling has been observed previously in the *mai-2* mutants [46]. Altogether, these experiments show that proper regulation of Δψ_m_ is required for the beneficial effects of DR in *C. elegans* and suggest a model whereby both ANT and IF1 activity play a role in the ability of DR to preserve Δψ_m_ during aging.

## DISCUSSION

In this study, we show that BD-induced DR in *C. elegans* attenuates the age-related loss of Δψ_m,_ and that this is necessary for lifespan extension and improved motility. The ability of DR to preserve Δψ_m_ during early aging does not appear to involve the activity of UCPs, but instead requires both mitochondrial ANT and IF1 activity. We also find that fatty acid metabolism is required for preservation of Δψ_m_ by DR, consistent with a recent report in worms that fatty acid metabolism promotes longevity [21], as well as numerous studies in mammals showing that DR increases fatty acid oxidation [36, 47-49]. While it remains difficult to quantify or control Δψ_m_ *in vivo* in mammals, this work should serve to orient future experiments to define mechanisms underlying these observations in worms and other systems. One limitation of our study is that UCP-4 in *C. elegans* may also regulate mitochondrial succinate transport in addition to its uncoupling role [40]. UCPs are closely related to metabolite transport proteins of many different varieties [50, 51] and different UCPs may even function differently within a single organism [52]. This is not a major limitation of our study because we found that *ucp-4* was not required for preservation of DR, Δψ_m_, and lifespan extension.

However, this caveat highlights the need for expanded understanding of UCP activity and DR in both invertebrate and mammalian systems. We do not, therefore, rule out a role for UCP activity in general for DR-mediated longevity in other models. On the contrary, investigating roles for UCPs under conditions of DR will be interesting given that UCPs are a popular target for diseases of aging, such as metabolic syndrome. There is evidence that targeting UCPs will be informative for understanding DR signaling in mammals, especially under conditions of protein restriction [51, 53, 54].

Our data supports a potential direct role for ANT and IF1 downstream of DR. In addition to regulating Δψ_m_, ANT and IF1 also modulate ATP/ADP dynamics in both mitochondria and cytosol, which may contribute to their effects on lifespan. ANT and IF1 seem to interact to modulate ATP synthase reversal under conditions of changing Δψ_m_ *in vivo* [44]. Our data suggest that this three-way interaction centered around Δψ_m_ could be important in the biology of DR. Developing tools to precisely control ATP/ADP levels in both the cytosol and in the mitochondrial matrix, as well as control over Δψ_m_, will be necessary first steps in elucidating more detail in experimental models. However, emerging evidence suggests that directly intervening on ANT function is possible to facilitate better understanding of mitochondrial energetics *in vivo* in the context of aging [18, 55]. Similarly, targeting IF1 in mammalian cells has been informative for probing the mechanisms of nutrient-sensing signaling and Δψ_m_ [56, 57].

Our results are also in line with our recent discovery that increased Δψ_m_ in itself is sufficient to extend lifespan in *C. elegans* [14]. We recently proposed an “energetics perspective on geroscience” [58]. The data presented here extend those ideas, showing that increased Δψ_m_ may be a fundamental mechanism of lifespan extension, at least in the case of DR for worms. Decreased Δψ_m_ has long been hypothesized to cause biological aging [13, 58-61] and the idea that improved Δψ_m_ with age is beneficial is not a new one [18, 58]. Now that this idea is established experimentally in worms, new perspectives on biological aging can be tested *in vivo*. It will be of particular interest to determine whether alternative interventions for increasing lifespan in worms and other organisms similarly impact Δψ_m._ In conclusion, we propose that *in vivo* Δψ_m_ is an important parameter during early phases of aging, and that longevity interventions to target Δψ_m_ will be informative for developing mitochondrial therapeutics for healthy aging.

## ACKNOWLEDGEMENTS

BJB is supported by the Biological Mechanisms for Healthy Aging (BMHA) Training Grant NIH T32AG066574. This work was supported by P30AG013280 to MK. Some strains were provided by the CGC, which is funded by NIH Office of Research Infrastructure Programs (P40 OD010440). The authors thank the laboratory of Rosa E. Navarro González for providing strain RN70 harboring the *mai-2* mutation.

