## Supplement for "Preservation of Mitochondrial Membrane Potential is Necessary for Lifespan Extension from Dietary Restriction"

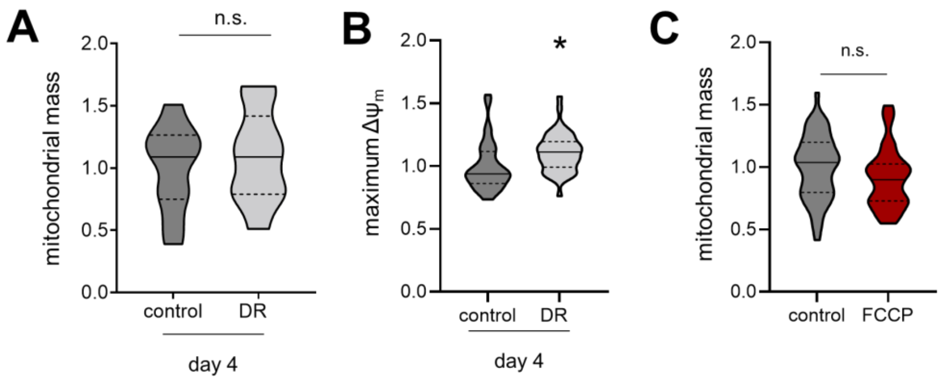


***Supplementary Figure 1. DR and FCCP effects on mitochondria in vivo.***

*A) Relative MitoTracker Green FM fluorescence in day 4 adult animals either fully fed or subjected to DR. Unpaired two-tailed t test, p = 0.5883. N = 10 animals each condition. Violin plots are medians ± quartiles (dotted lines). B) Maximum TMRE fluorescence in day 4 adult animals either fully fed or subjected to DR. Unpaired two-tailed t test, *p = 0.0070. Control N = 34, DR N = 44. Violin plots are medians ± quartiles (dotted lines). C) Relative MitoTracker Green FM fluorescence in day 4 adult animals treated with either vehicle control or 10 µM FCCP. Unpaired two-tailed t test, p = 0.1380. Control N = 52, FCCP N = 32. Violin plots are medians ± quartiles (dotted lines).*

***Supplementary Table 1. Lifespan analyses.*** Statistical information to support lifespans in each indicated figure. All experiments were carried out in the same environment. For each panel related to Figure 3, wildtype comparisons are made to the control N2 lifespan from Figure 2B, which is highlighted with gray text in the table.

| Figure | Strain | Treatment | Median lifespan (days) | compared to: | comparison p value (log-rank (Mantel-Cox) test) | # of animals |
|---|---|---|---|---|---|---|
| Fig2B | N2 | control | 19 | | | 161 |
| | N2 | DR | 24 | control | < 0.0001 | 116 |
| | N2 | FCCP | 18 | control | 0.14 | 158 |
| | N2 | DR+FCCP | 17 | DR | < 0.0001 | 90 |
| Fig2C | N2 | control | 20 | | | 95 |
| | N2 | FCCP | 20.5 | N2 control | 0.3 | 88 |
| | *eat-2* | control | 25 | N2 control | < 0.0001 | 159 |
| | *eat-2* | FCCP | 22 | *eat-2* control | < 0.0001 | 216 |
| Fig3F | *ucp-4* | control | 16 | | | 257 |
| | *ucp-4* | DR | 24 | control | < 0.0001 | 73 |
| | *ucp-4* | FCCP | 16 | control | 0.95 | 164 |
| | *ucp-4* | FCCP+DR | 16.5 | DR | < 0.0001 | 92 |
| | *ucp-4* | control | 16 | N2 control | < 0.0001 | 257 |
| Fig3G | *ant-1.2* | control | 22 | | | 186 |
| | *ant-1.2* | DR | 24 | control | 0.13 | 165 |
| | *ant-1.2* | FCCP | 19 | control | < 0.0001 | 164 |
| | *ant-1.2* | FCCP+DR | 19 | DR | < 0.0001 | 172 |
| | *ant-1.2* | control | 22 | N2 control | <0.0001 | 186 |
| Fig3H | *mai-2* | control | 23 | | | 139 |
| | *mai-2* | DR | 24 | control | 0.0072 | 139 |
| | *mai-2* | FCCP | 18 | control | < 0.0001 | 112 |
| | *mai-2* | FCCP+DR | 21 | DR | 0.09 | 126 |
| | *mai-2* | control | 23 | N2 control | <0.0001 | 139 |
